# Associative Learning from Replayed Experience

**DOI:** 10.1101/100800

**Authors:** Elliot A. Ludvig, Mahdieh S. Mirian, E. James Kehoe, Richard S. Sutton

**Author notes:** Corresponding Author: Elliot A. Ludvigs, Department of Psychology, University of Warwick Coventry, UK, CV4 7AL.

## Abstract

We develop an extension of the Rescorla-Wagner model of associative learning. In addition to learning from the current trial, the new model supposes that animals store and replay previous trials, learning from the replayed trials using the same learning rule. This simple idea provides a unified explanation for diverse phenomena that have proved challenging to earlier associative models, including spontaneous recovery, latent inhibition, retrospective revaluation, and trial spacing effects. For example, spontaneous recovery is explained by supposing that the animal replays its previous trials during the interval between extinction and test. These include earlier acquisition trials as well as recent extinction trials, and thus there is a gradual re-acquisition of the conditioned response. We present simulation results for the simplest version of this replay idea, where the trial memory is assumed empty at the beginning of an experiment, all experienced trials are stored and none removed, and sampling from the memory is performed at random. Even this minimal replay model is able to explain the challenging phenomena, illustrating the explanatory power of an associative model enhanced by learning from remembered as well as real experiences.

## Associative Learning from Replayed Experience

Associative models provide powerful explanations for many learning phenomena (e.g., Bush & Mosteller, 1951; Enquist et al., 2016; Harris, 2006; Ludvig et al., 2011, 2012; Miller & Shettleworth, 2007; Pearce & Bouton, 2001; Pearce & Hall, 1980; Rescorla & Wagner, 1972; Sutton & Barto, 1981). Many of these models use an error-correction learning rule, whereby learning is driven by the difference between what was expected and what actually occurred. This difference drives a change in associative strength that determines later performance. In this paper, we propose that the same error-correction rule for learning from externally generated experience can also be used on replayed experiences retrieved from memory. This extended rule allows learning from previously encountered events stored in memory, thus allowing associative strengths to change without further direct experience. This simple idea leads to a model, termed the *replay model*, which provides a unifying explanation for important learning phenomena that have proved challenging to earlier associative models, including spontaneous recovery, latent inhibition, retrospective revaluation, and trial spacing effects.

The replay model is formally an extension of the Rescorla-Wagner (RW) model (Rescorla & Wagner, 1972), which has been the dominant account of associative learning for more than 40 years (Ludvig et al., 2011; Miller et al., 1996). Despite numerous successes, the RW model has significant limitations in its scope (Miller et al., 1996). For example, one difficulty for the RW model is the phenomenon of spontaneous recovery (Haberlandt et al., 1978; Napier et al., 1992; Pavlov, 1927; Rescorla, 2004). In spontaneous recovery, a conditioned response that has been extinguished will gradually reappear with the passage of time. The RW model cannot explain this change in responding as the model only allows learning about events at the time they are encountered in the real world.

The idea that animals replay past experiences has been used in neuroscience to explain the behaviour of place cells in the hippocampus (e.g., Davidson et al., 2009; Euston et al., 2007; Gupta et al., 2010; Redish, 2016; Wilson & McNaughton, 1994). When animals navigate through an environment, place cells become active in a sequence corresponding to their preferred spatial location. Following such a task, these neurons exhibit experience replay: The same correlated patterns of activity occur after the spatial task as during the task, mostly during short-wave sleep, but also during awake post-task periods. For example, Gupta et al. (2010) trained rats to find food in one of two side locations in a maze shaped like a figure eight. They recorded from hippocampal place cells in resting rats and found several distinct patterns of place-cell reactivations. The place cells replayed recent and remote trajectories of the rat in both forward and backward directions. Similar patterns of reactivation are also observed in zebra finches following song learning (e.g., Dave & Margolish, 2000). In human memory tasks, item-specific reactivations are also sometimes observed, which correlate with later recall (Deuker et al., 2013; Staresina et al., 2013). This variety of replay patterns suggests that the brain directly rehearses both recent and remote past experiences.

Re-using past experience to enhance associative processes has achieved some success in artificial intelligence, especially in reinforcement learning. For example, Lin (1992) showed how replay of previous experience could accelerate learning performance for a reinforcement-learning agent in a complex grid world. Recently, a deep learning network enhanced with experience replay achieved human-level performance on a suite of Atari video games (Mnih et al., 2015). More generally, this past experience can be abstracted into a model of the world for the agent, which can then be used to generate trajectories of imagined experience. The *Dyna* class of model-based reinforcement-learning algorithms follows this approach, by using the same associative algorithm to learn from both model-generated trajectories and real experience (Shohamy & Daw, 2015; Johnson & Redish, 2005; Silver et al., 2008; Sutton, 1990; Sutton et al., 2008; van Seijen & Sutton, 2015). Similar model-based reinforcement-learning approaches have also been used in the context of conditioning to explain differences between habits and goal-directed behavior (Balleine & O’Doherty, 2009; Daw, Niv & Dayan, 2005; Daw et al., 2011; Dolan & Dayan, 2013; Miller, Shenhav, & Ludvig, 2016). The replay model also combines learning from internally generated experiences with learning from external trials, but those internally generated experiences are strictly replays of past trials. Because the replay model operates at a trial level, there is no abstract model of the world to be learned, making remembered and imagined experience equivalent in this case.

The replay model builds on two familiar concepts in memory research: consolidation and rehearsal, which have been used to describe processes that underpin long-term storage of recent events (e.g., Atkinson & Shiffrin, 1968; Chapman, 1991; McClelland et al., 1995; McGaugh, 2000; Ratcliff, 1990, Stickgold, 2005). Consolidation refers to the strengthening of memory traces that sometimes occurs after initial exposure, which may occur through the explicit revisiting or rehearsal of past experience. As used here, the concept of “replay” represents an enhanced rehearsal-like process—one which engages an error-correction learning rule in the same way as external events. This combination provides a mechanism through which rumination about remote past experiences plays a key role in the process of associative learning and may lead to the apparent consolidation of memories.

Rehearsal processes have previously been proposed to play a limited role in associative learning. For example, Wagner et al. (1973) showed that inserting a surprising post-trial experience during classical eyelid conditioning in rabbits slowed conditioning to the target stimulus, concluding that this additional experience interfered with rehearsal. These results suggested that rehearsal may preferentially occur to surprising outcomes (cf. Salafia & Papsdorf, 1968; Salafia et al., 1977). Rehearsal processes have also been used to explain serial order effects in human associative learning (e.g., Chapman, 1991; Gershman et al., 2013; Melchers et al., 2004; Ratcliff, 1990). Here we develop the replay model, which implements these ideas more comprehensively and connects them directly to error-correction. We then explore the implications of this model over a range of classical conditioning experiments, showing how the model provides new insights into some previously intractable aspects of associative learning.

### Model Details

For purposes of simulation, we adapted the competitive, error-correction rule as originally implemented in the Rescorla-Wagner (RW) model. The RW rule works by calculating the net associative strength (*V^Σ^*) on each trial *T* as the sum of the associative strength for all stimuli available on that trial:

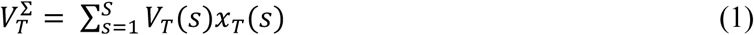

where *V* represents the individual associative strengths for each stimulus s, and *S* is the total number of stimuli possible in the experiment. The indicator variable x represents whether or not a particular stimulus was present on a given trial and is set to 1 when present and 0 when absent. This net associative strength is then compared against the actual US or reward *r* to generate a prediction error *δ* for that trial:

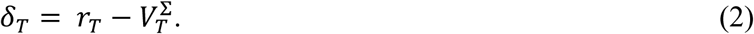

The associative strengths for each of the stimuli present on that trial are then updated with a fixed proportion (*α*) of this prediction error:

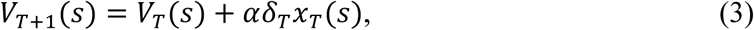

where *x* is again the same indicator variable as above. Through this learning process, the net associative strength *V* becomes a prediction of the total reward on the trial. The learning-rate parameter *α* is usually confined to be between 0 and 1.

The replay model uses this same learning rule for updating associative strengths, but deploys the learning rule in a wider range of situations than the original RW formulation. As depicted in Figure 1, there are two parallel streams of information processing in the replay model. First, there is the usual process of associative learning as formalized in the original RW model (direct pathway): The animal encounters the stimuli and rewards on a trial and updates the relevant associative strengths in the same manner as the RW model. Second, in the indirect, replay pathway, the events in a trial accumulate in a trial memory. The trial memory thus contains an event record (stimuli, rewards) for past trials. Samples from this trial memory are continuously drawn and replayed (both during trials and in between trials). In this replay process, a new prediction error is calculated based on the current associative strengths, and the relevant associative strengths are updated using the RW error-correction rule (Eqs. 2-3).

**Figure 1.**
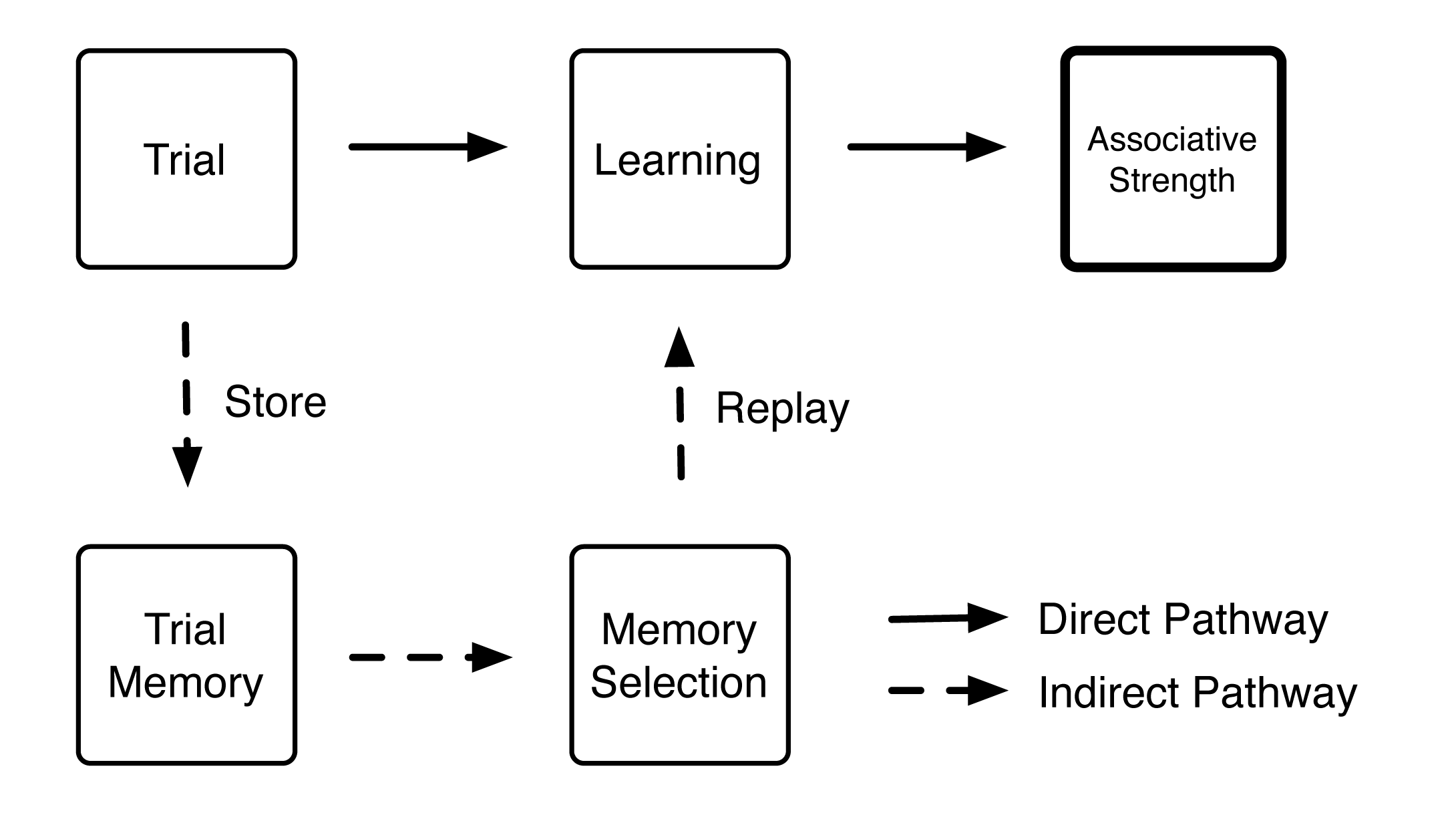
Schematic of the replay model. There are two processing pathways in the model. First, the model updates its associative strength by comparing the predicted reward with the actual outcome on that trial (solid arrows). In addition, trials are stored in a trial memory in the indirect pathway and are then sampled and replayed (dashed arrows). A replayed trial is treated like any other: The model updates its associative strength by comparing the predicted reward with the actual outcome on the replayed trial.

To demonstrate the full explanatory power of the model, we simulated four disparate, yet well-established, learning phenomena. They are spontaneous recovery, latent inhibition, retrospective revaluation, and trial spacing effects. These four phenomena are robust features of associative learning, which have been replicated many times in different species and different preparations (e.g., Schmajuk & Alonso, 2012). Our strategy is to focus on qualitative simulation of the core phenomena; detailed fitting to the exact parameters of particular datasets is left to future work. Previously, each of these four phenomena has had a tailored explanation within an associative framework, but no single theory has been able to explain all of them. The sections below will each include a description of the basic phenomenon, a brief description of the existing associative explanations, and the simulation results from the replay model.

For the simulations, we used a minimal version of the replay model, with a simple set of assumptions about how memories are stored and sampled. To start each simulation, the trial memory was assumed to be empty. Over the course of learning, representations of all experienced trials were progressively accumulated in the trial memory, and none were ever removed. For each replay, the trial to be replayed was randomly selected from that full set of experienced trials. As a result, the likelihood that a particular type of trial was sampled for replay was proportional to the frequency with which it had occurred previously. Below we consider some implications of relaxing these assumptions, allowing for a variety of storage, curation, and selection schemes. The learning-rate parameter (*α*) was set to .05 for real trials and to .005 for replayed trials, and there were 7 replay periods between each external trial.

### Model Mechanics: Simple Acquisition

To illustrate the basic mechanics of the model, Figure 2 shows how the model works during the simple acquisition of a classically conditioned response. During simple acquisition, a stimulus is repeatedly paired with a reward. For example, a tone might be repeatedly paired with food or a light might be repeatedly paired with a puff of air to the eye. Animals generally learn to predict the relevant reward (food or puff of air) and make an appropriate response (e.g., salivate or blink). For this simulation, the conditioned stimulus X was simply paired with reward on four trials, indicated by the symbol X+ (top row). On each trial, two things happen: first, the trial events (stimulus and reward: X+) are stored in the trial memory, which grows with each trial (middle row). Second, the associative strength is incremented using the RW learning rule (Eq 1; bottom row). Between trials is where the replay model becomes distinct from the basic RW model: Acquisition trials (X+) are sampled from the trial memory and replayed, leading to further increments in the associative strength. In this simple-acquisition scenario, the additional processing leads to faster learning of the association between X and the reward. As may be apparent, for this simple-acquisition scenario, the additional processing would be invisible. As will be shown in the next sections, however, the impact of the additional processing becomes highly visible as more variables are manipulated.

### Spontaneous Recovery

Spontaneous recovery has proven to be particularly difficult to reconcile with the RW model and similar associative accounts of learning (Bouton, 1993; Pavlov, 1927; Kehoe, 1988; Rescorla, 2004; Sissons & Miller, 2009). In a spontaneous recovery experiment, acquisition training using reinforced presentations of a conditioned stimulus (as above) is followed by extinction training containing stimulus-alone presentations. During the acquisition training, animals learn to respond to the stimulus (see Fig 2), but then they progressively cease responding during extinction training. If, however, after the end of extinction training, the stimulus is represented to the animals after a delay, the extinguished response reappears, sometimes at nearly full strength (Kehoe, 2006; Napier et al., 1992). The degree of recovery increases as the delay between the end of extinction training and recovery testing is increased (e.g., Haberlandt et al., 1978—see Figure 3C).

**Figure 2.**
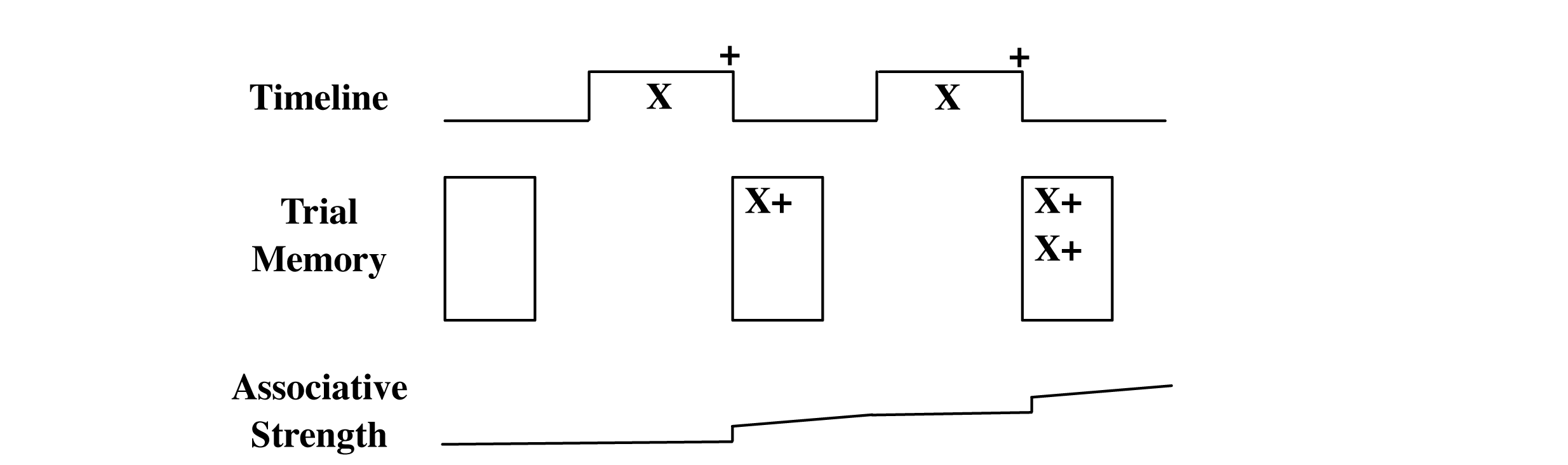
Illustrative example of the replay model during simple acquisition. The top row represents the experimental sequence, where X represents the stimulus and + represents the reward. The middle row represents the trial memory, which grows with each successive trial. The bottom row depicts the growth in associative strength across conditioning, both due to the learning from the real trials and from the replayed trials during the inter-trial intervals.

**Figure 3.**
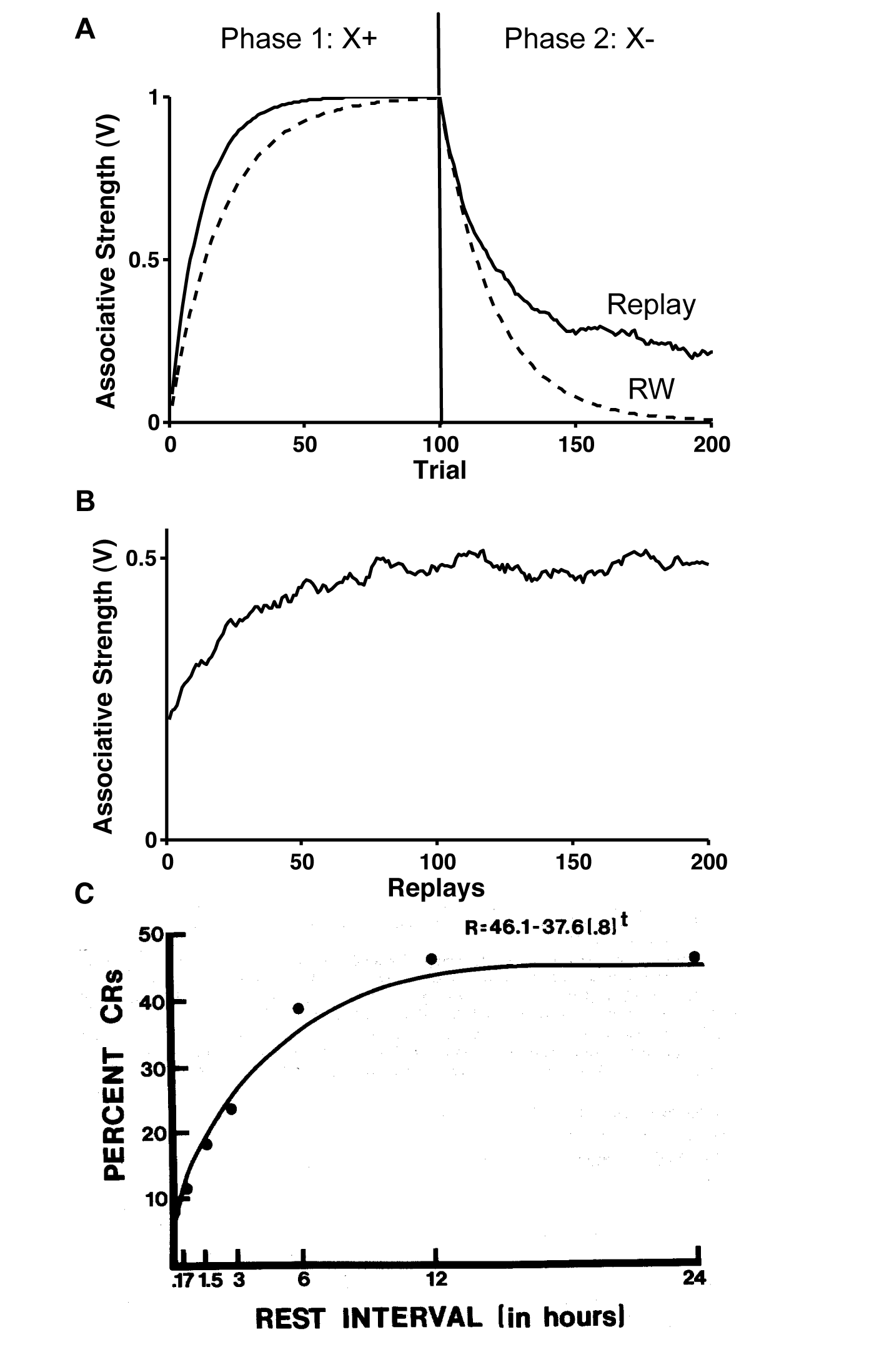
Spontaneous Recovery. (A) Simulation results from the replay model (solid line) and the Rescorla-Wagner (RW) model (dashed line) during the initial acquisition (X+) and extinction phases (X-). (B) Associative strength of X as a function of the number of replays since the end of the extinction phase in the replay model. Note the change in the scale of the x-axis. (C) Data from Haberlandt et al. (1978) showing the degree of spontaneous recovery after different post-extinction delays in rabbit eyeblink conditioning (permission to reproduce pending).

According to the RW model, at the end of extinction, the associative strength (V) approaches 0. There is no mechanism for any change in this associative strength during the intervening interval before the recovery test. Thus, the original RW model cannot account for spontaneous recovery. Within the wider associative framework, numerous explanations for spontaneous recovery have been suggested, including different decay rates for excitatory and inhibitory processes (Bouton, 1993; Pan et al., 2008), different decay rates for the first and second things learned (Bouton, 1993; Devenport, 1998; Sissons & Miller, 2009), changes in the sampled stimulus characteristics over time (Estes, 1955), and the inference of different underlying states of the world for acquisition and extinction (Gershman, Blei & Niv, 2010; Gershman & Niv, 2012).

The replay model augments the RW model by assuming that during the delay between the end of extinction and the recovery test, both acquisition trials and extinction trials stored in the trial memory are replayed randomly. As a result of the replays of the acquisition trials, the associative strength recovers to an intermediate value. Figure 3 shows a simulation of spontaneous recovery. In the simulation, the replay model was first trained with 100 acquisition trials (X+) followed by 100 extinction trials (X−). Associative strength increases to an asymptote near 1 during acquisition and then decreases toward 0 during extinction (Fig 3A).

At the end of the extinction phase, the model is not given any further training trials, but is allowed to continue replaying previously experienced trials. In this simplest formulation of the replay model, the trials replayed only depend on the frequency with which those trial types have appeared in the past. Because both acquisition trials and extinction trials appeared equally often in the initial training, with further replay (i.e., more time), the associative strength of stimulus X approaches 0.5 (Fig. 3B). This pattern resembles the degree of recovery exhibited by the conditioned eyeblink response in rabbits as the delay between extinction and the recovery test is increased (Haberlandt et al., 1978; Fig 3C). The replay model makes some clearly testable predictions: for example, the degree of spontaneous recovery should depend on the relative number of acquisition and extinction trials. With an increased number of extinction trials, recovery should be muted, as is observed in fear conditioning (e.g., Laborda & Miller 2013).

### Latent Inhibition

Latent inhibition occurs when animals are initially exposed to unreinforced presentations of a stimulus. These initial exposures reduce the speed of conditioning when the stimulus is later paired with a reward (Lubow, 1973; Lubow & Moore, 1959). This finding poses a challenge to the original RW model, because the RW model does not produce changes in associative strength during the initial stimulus exposures when there is no expected reward and thus no prediction errors occur. Other prominent associative learning models have postulated an attentional mechanism, whereby animals learn not to attend to the stimulus when it is repeatedly presented alone during the first phase of the experiment (Mackintosh, 1974; Pearce & Hall, 1980). The replay model explains latent inhibition by supposing that during acquisition training, when the animal receives pairings of the stimulus with a reward, the animal also replays some of the earlier unreinforced, stimulus-only trials, thereby retarding conditioning.

Figure 4 presents results from simulations of the replay model (top) and the original RW model (bottom) with a latent inhibition procedure. In the first phase, the models are presented initially with either 100 unreinforced presentations of the stimulus that will later be conditioned (X-, solid line) or a different stimulus (Y-, dashed line). In the second phase for both conditions, the models are presented with 100 reinforced presentations of the target stimulus (X+). As with real animals, neither the replay model nor the RW model show any change in associative strength during the stimulus-alone presentations in the first phase. In the second phase, however, a difference between the two models emerges. In the replay model (Fig. 4A), the acquisition to the stimulus depends on its training history, whereas, in the RW model, acquisition proceeds equivalently in both groups. As shown in Figure 4A, if the replay model is exposed to X- and then X+, then the replay of X- trials during the second phase slows down acquisition. This replay of unreinforced X- trials acts like intercalated extinction trials, slightly decrementing the associative strength of the pre-exposed stimulus (X). In contrast, when the model is exposed to an unrelated stimulus (Y-) and then X+, then the replay of the Y- trials during X+ training has no direct effect on the associative strength of X, and conditioning proceeds as though no preexposure had occurred.

**Figure 4.**
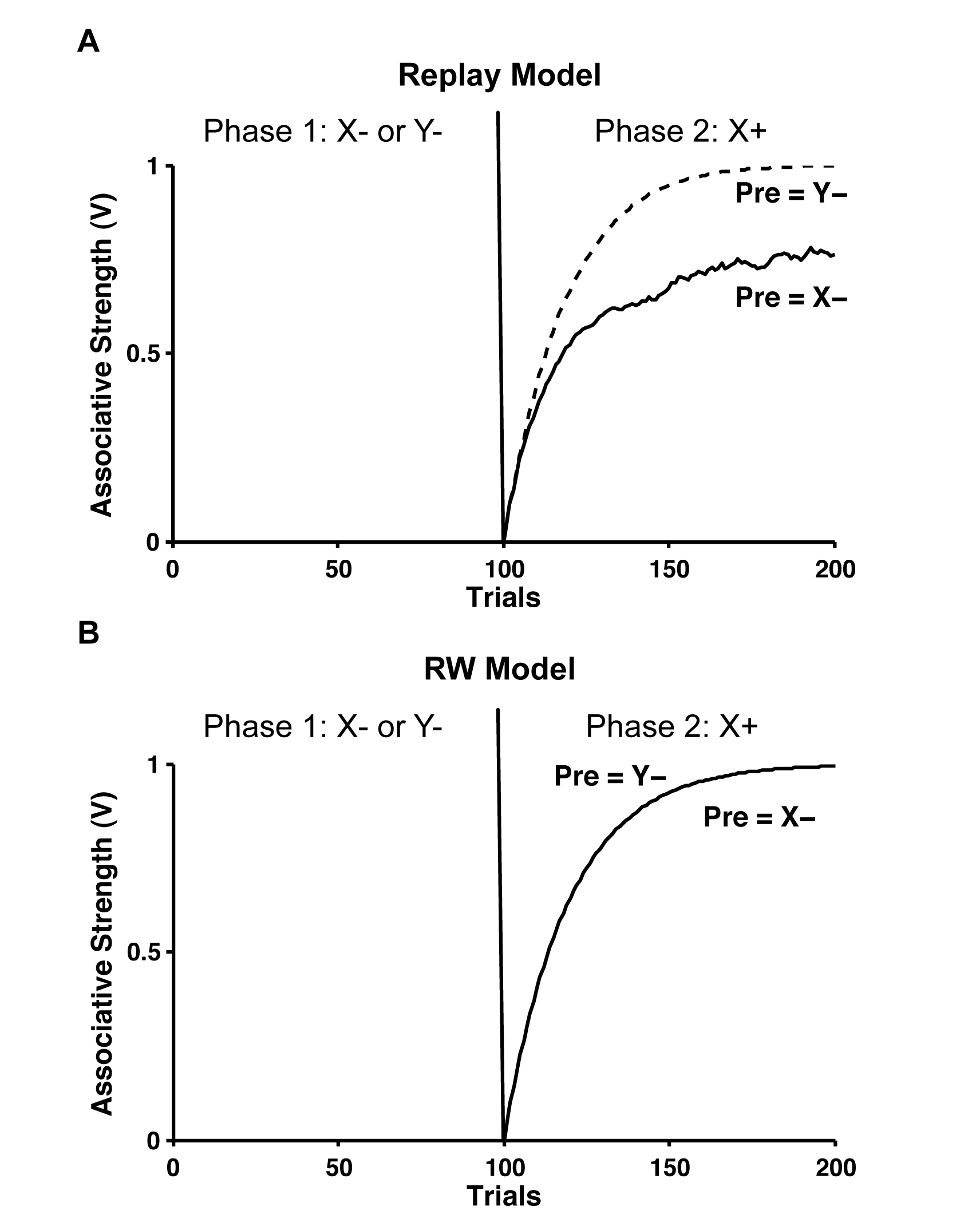
Simulation results in a latent inhibition experiment. (A) In the replay model, with pre-exposure to the same stimulus (Pre = X-; solid line), conditioning proceeds more slowly during acquisition (X+) than comparable pre-exposure to a different stimulus (Pre = Y-; dashed line). Both lines depict the associative strength of X (i.e., V(X)). (B) In the Rescorla-Wagner (RW) model, there is no difference between the two pre-exposure groups during acquisition training; accordingly, both lines in the bottom panel completely overlie.

### Retrospective Revaluation

A third class of associative learning phenomena that can be reproduced by the replay model is retrospective revaluation (e.g., Dickinson & Burke, 1995; Van Hamme & Wasserman, 1994). In retrospective revaluation, animals change their responding to one stimulus as a result of further training with a different stimulus. For example, in backwards blocking (Miller & Matute, 1996; Shanks, 1985), two stimuli are initially paired with a reward (XY+) followed by pairing of only one of those two stimuli with the same reward (X+). Animals initially learn to respond to both stimuli, but will cease responding to the absent stimulus (Y) if tested after the second phase of training (X+) despite never re-encountering the Y stimulus (see Figure 5).

Retrospective revaluation has been explained by a combination of within-compound associations plus negative learning rates associated with absent, but associatively-activated, stimuli (e.g., Dickinson & Burke, 1995; Van Hamme & Wasserman, 1994). According to these explanations, a representation of the absent stimulus (Y) is activated by the presented stimulus (X), and this representation has a negative learning rate (*α* < 0). As a result, presentation of the reinforced trial (X+) leads to a decrement in the associative strength of the absent stimulus (Y). Other explanations of retrospective revaluation have included Bayesian belief updating based on the new information (Daw & Courville, 2008), an interaction between overlapping stimulus representations (Ghirlanda, 2005), and instance-based memory recall of the absent stimulus (Jamieson et al., 2010).

The replay model offers a different explanation. During the latter phases of training, trials from earlier phases are sampled from the trial memory and replayed, updating the associative strengths through the same learning rules as for the initial acquisition training (see Chapman, 1991, and Melcher et al., 2004 for related explanations). To test the scope of our replay model in explaining retrospective revaluation, we considered three examples: backwards blocking, recovery after blocking, and backwards conditioned inhibition.

### Backwards Blocking

As described above, in backwards blocking, animals are first presented with a reinforced compound stimulus (XY+) and then are trained with one of those stimuli reinforced in isolation (X+). When tested after this second phase, animals decrease responding to the other stimulus (Y) that was not presented in the second phase, provided that the reward has low biological significance (Shanks, 1985; Miller & Matute, 1996). Figure 5 shows simulations from the replay model and the original RW model in a backwards-blocking experiment. In the first phase, the models are presented with 100 reinforced compound-stimulus presentations (XY+), and the associative strength of each element (X, Y) of the compound grows to 0.5. In the second phase, the models are trained with 100 additional X+ trials. In the replay model, the associative strength for reinforced stimulus X continues to increase, while the associative strength of the absent stimulus Y decreases (Fig. 5A). According to the model, this decrement in the associative strength occurs because the compound XY+ trials continue to be replayed during X+ training. Because the associative strength of X+ has continued to grow, during these XY+ trials the net associative strength is greater than the total reward, producing overexpectation and a decrement in the associative strength of both X and Y. For stimulus X, this decrement is offset by the increment from the ongoing X+ trials, whereas, for the absent stimulus Y, there are only decrements. In contrast, in the original RW model (Fig. 5B), only the associative strength of the presented stimulus X is changed in the second phase of the experiment.

**Figure 5.**
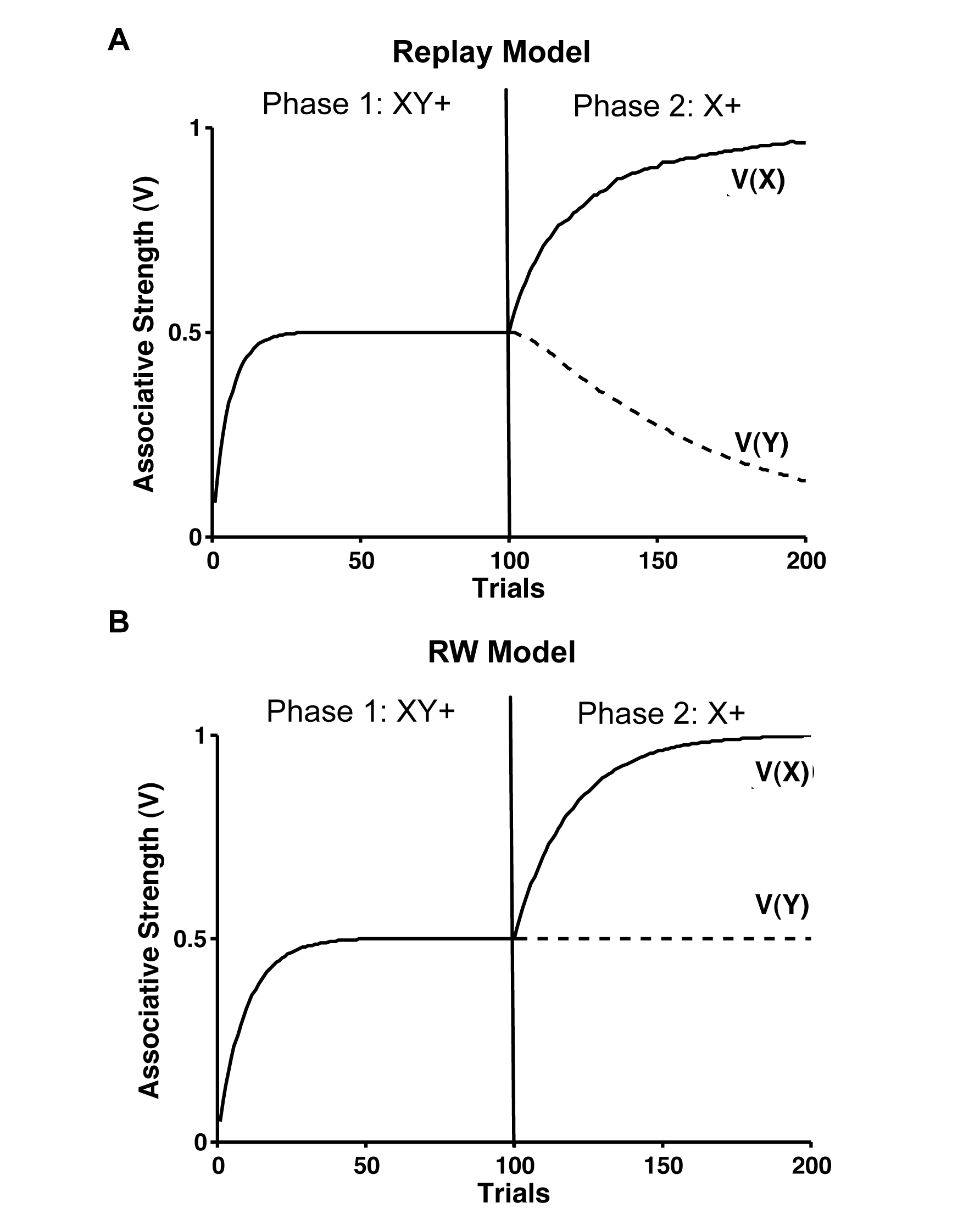
Backwards Blocking. In backwards blocking, compound-stimulus training (XY+) is followed by reinforced presentation of one of the elements (X+). (A) In the replay model, both stimuli initially gain associative strength equally, asympoting at 0.5. Once the elemental training begins, the associative strength of the reinforced element continues to grow, whereas the associative strength of the no-longer-presented stimulus gradually decreases. (B) The Rescorla-Wagner (RW) model behaves similarly in the first phase (XY+), but there is no change in the associative strength of the unpresented stimulus during the second phase (Y+).

### Recovery after Blocking

In a recovery after blocking experiment, animals are first trained with a standard blocking protocol (X+ then XY+) followed by extinction of the blocking stimulus (X-). At the end of the standard blocking protocol, animals show limited responding to the added stimulus (Y). During the X- extinction training, however, the level of responding to stimulus Y increases (e.g., Blaisdell et al., 1999). Figure 6 shows how the replay model and original RW model perform on the three phases of this experiment. During the initial X+ training, the associative strength of stimulus X increases to asymptote in both models, with no change in the strength of the yet-to-be-experienced stimulus (Y). In the second phase, during XY+ training, there are no prediction errors and thus no learning, as the reward is still fully predicted by stimulus X. The additional replays of X+ or XY+ do not alter the associative strengths—maintaining the blocking effect. In the final X- extinction phase, the associative strength of stimulus X declines in both models. In the original RW model, this decrement is the only change in associative strength; Y’s associative strength remains at 0 (Fig. 6B). In the replay model (Fig. 6A), however, previous trials are replayed, including the XY+ trials, resulting in an increase in the associative strength of the previously blocked stimulus (Y). This increase is due to positive prediction errors on the replayed XY+ trials, which are caused by the decrease in the associative strength of stimulus X.

**Figure 6.**
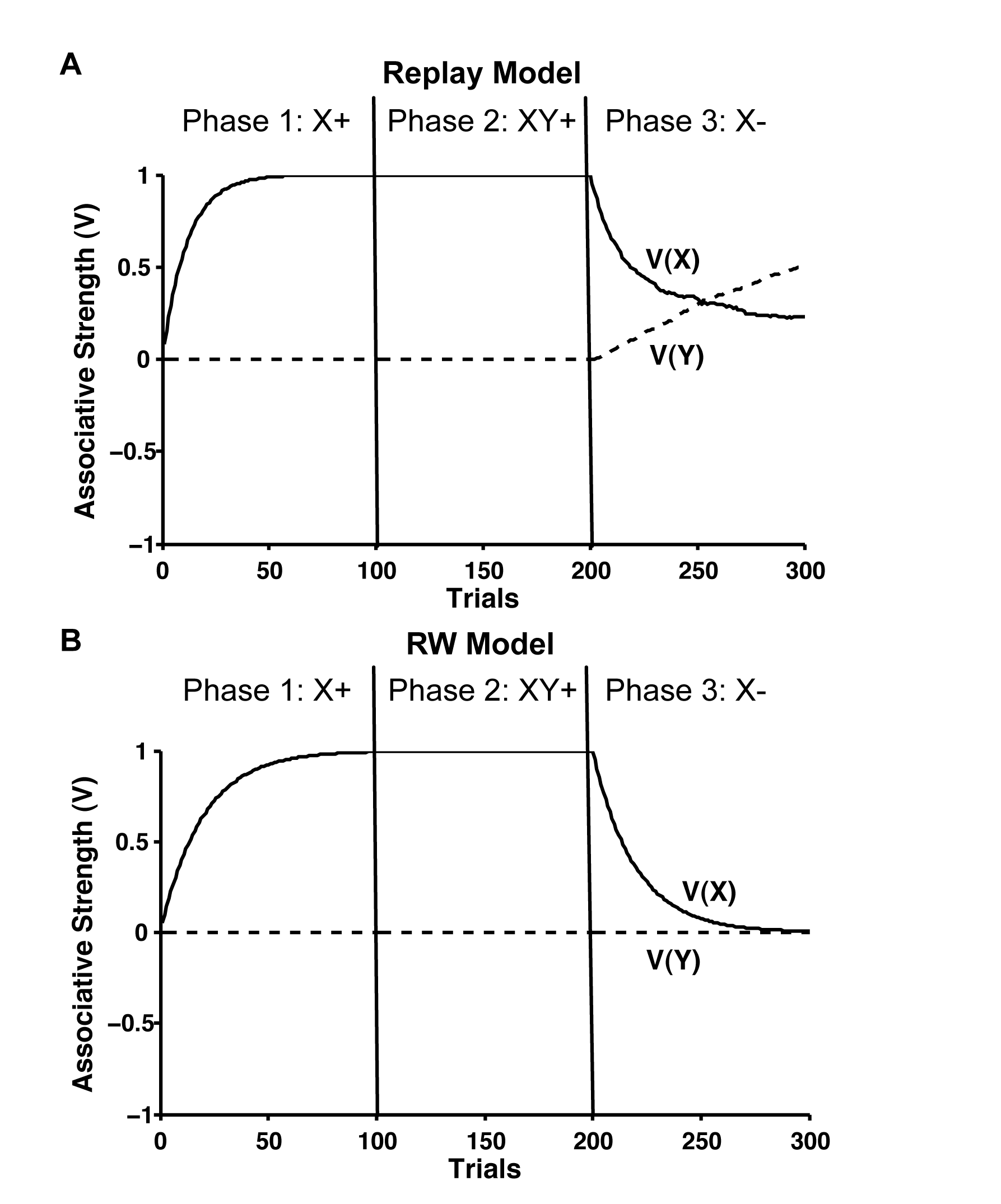
Recovery after Blocking. (A) Simulated results from the replay model show how during the initial acquisition training (X+) and subsequent blocking training (XY+), the associative strength (V(X)) of the blocking stimulus X reaches an asymptote of 1, while the associative strength (V(Y)) of the blocked stimulus stays at 0. In the final recovery phase (X-), the associative strength of stimulus X decreases as expected, but the associative strength of stimulus Y grows because of occasional replays of the compound trials (XY+) from the blocking phase. (B) The Rescorla-Wagner (RW) model shows similar results during the first two phases, but there is no increase in the associative strength of the unpresented stimulus Y during the last phase.

### Backward Conditioned Inhibition

As a third example of retrospective revaluation, we consider a situation that has been primarily demonstrated in human contingency learning: backwards conditioned inhibition (e.g., Chapman, 1991; Urcelay et al., 2008). This procedure adds an interesting twist to the other types of retrospective revaluation in that an absent stimulus becomes inhibitory. In a backwards conditioned inhibition procedure, the first phase of training consists of unreinforced presentations of a stimulus compound (XY-). In the second phase of training, one of the stimuli is reinforced (X+), and the other stimulus (Y) is no longer presented. The reinforced stimulus (X) becomes a conditioned exciter, whereas the no-longer-presented stimulus (Y) becomes a conditioned inhibitor, acquiring negative associative strength.

Figure 7 shows how the replay model can explain the results in backwards conditioned inhibition. There is no change in associative strength during the initial XY- training, but the associative strength for these two stimuli go in opposite directions during the subsequent X+ training: Positive associative strength accrues to the reinforced stimulus (X), while negative associative strength accrues to the unreinforced stimulus (Y), making it a conditioned inhibitor. This latter effect occurs in the replay model because the initial unreinforced compound trials (XY-) are replayed during the second phase, but now, the expected reward is greater than 0 due to the associative strength of the reinforced stimulus X. These unreinforced trials thus generate a negative prediction error, converting Y into a conditioned inhibitor. The original RW model cannot account for this result because stimulus Y is not re-presented during the X+ training phase (see Fig 7B).

**Figure 7.**
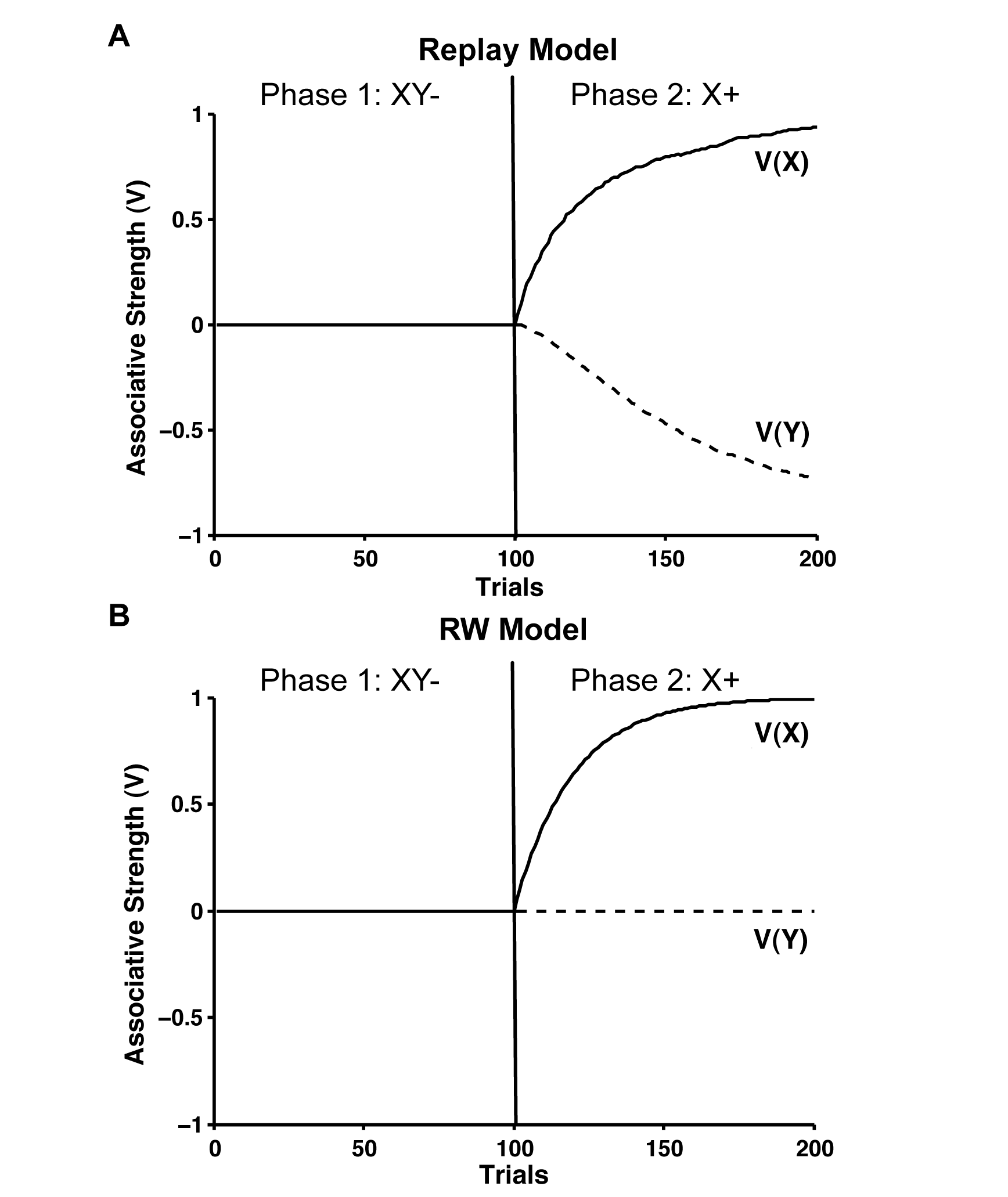
Backwards conditioned inhibition. (A) Simulated results from the replay model show the changes in associative strength (V) during a backwards conditioned inhibition experiment. In the first phase, there are unreinforced presentations of both stimuli together (XY-), and there are no changes in associative strength. In the second phase, only one stimulus is reinforced (X+), and the associative strength of that stimulus grows, while the associative strength of the second stimulus becomes negative (inhibitory) due to replays of the initial unreinforced compound trials (XY-). (B) In the Rescorla-Wagner (RW) model, there are no changes in the associative strength of unpresented stimulus Y during the final phase, and that stimulus does not become a conditioned inhibitor.

In all three of these simulations of retrospective revaluation, the overall qualitative result is not parameter dependent, but the degree of change in associative strength due to the retrospective revaluation depends on the two model parameters that control the strength of the replay relative to the initial experience. A higher learning rate (a) or greater number of replays per trial would result in more retrospective revaluation of the absent stimulus.

### Trial Spacing Effects

In addition to the above experiments that involve changes in the contingencies of reinforcement, the replay model also readily addresses some temporal features of associative learning, most notably trial-spacing effects, including timescale invariance. As in many other domains of learning (e.g., Brown, Roediger, & McDaniel, 2014), trials which are spaced further apart in classical conditioning yield faster learning than trials which are massed together (Barnet et al., 1995; Gallistel & Gibbon, 2000; Kehoe et al., 1991; Pavlov, 1927). This finding follows naturally from the core assumptions of the replay model (see Fig 2): during simple acquisition, spaced trials with longer intertrial intervals (ITIs) would allow for more replays than massed trials, thereby producing faster acquisition on a trial-by-trial basis. On an even longer timescale, the overnight increases in conditioned responding sometimes observed (e.g., Kehoe & White, 2002) is similarly explicable by assuming occasional replay of acquisition trials even outside the experimental session.

Timescale invariance is a stronger version of these basic trial-spacing effects and occurs when the rate of conditioning is mostly determined by the ratio between the duration of a trial and the duration of the inter-trial interval (see Balsam et al., 2010; Gallistel & Gibbon, 2000; 2002; Fig 8B). The RW model and other trial-level associative models that take little account of the ITI are strongly challenged by the finding of timescale invariance in conditioning (e.g., Gallistel & King, 2009). To explain time-scale invariance, the duration of the stimulus on reinforced trials must be included in the computations. Accordingly, we added the assumption that the duration of a replayed trial is proportional to the duration of the actual trial—an assumption that has some support in the replay rate of spatial experience (Davidson et al., 2009). As a result, longer trials lead to less-frequent replay and slower learning. With this extra assumption, timescale invariance in the rate of conditioning follows straightforwardly in our replay model.

**Figure 8.**
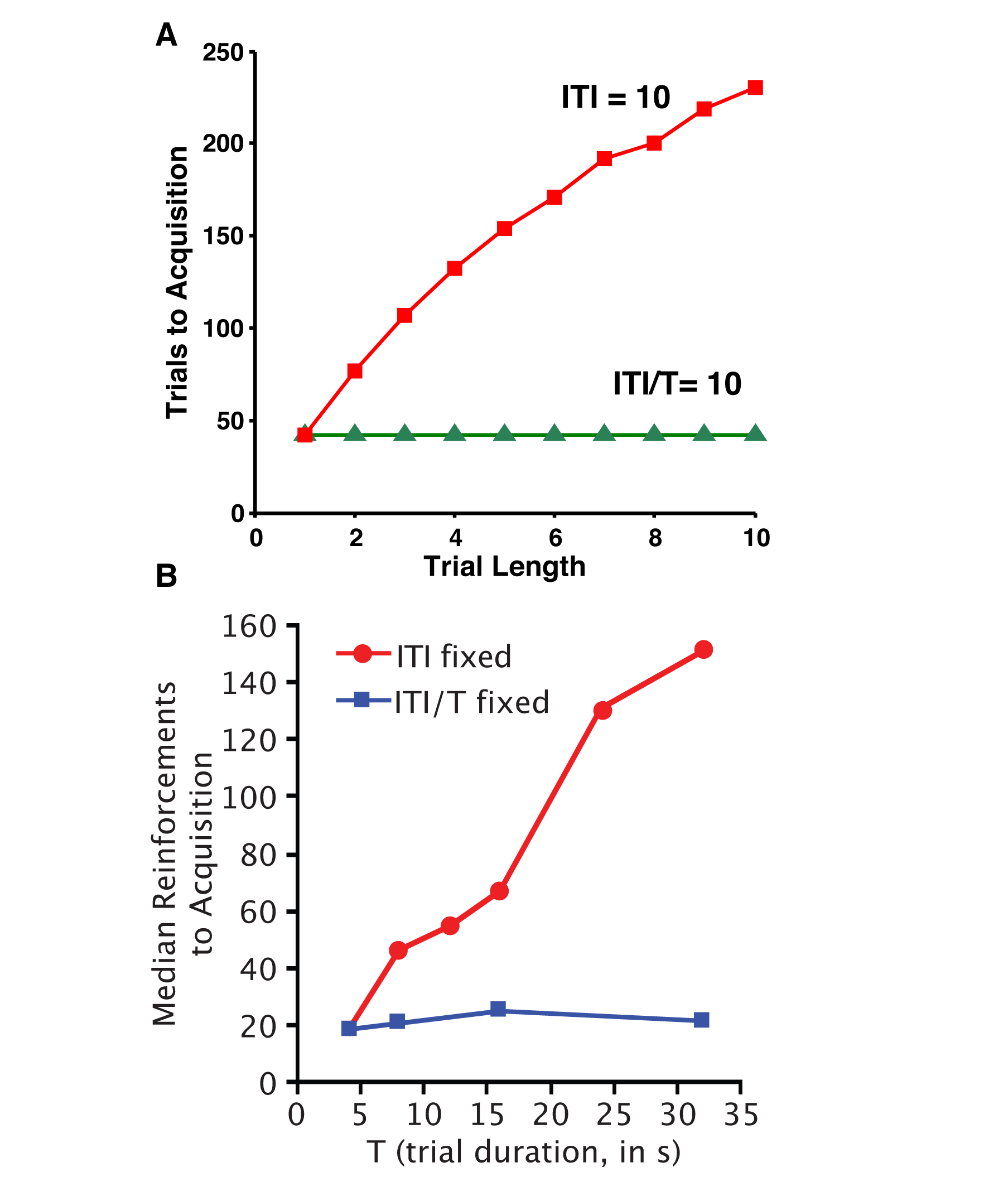
Timescale invariance. (A) Simulations from the replay model with replay speed inversely proportional to the trial duration—the longer the trial the more time it takes to replay. A constant ITI results in faster learning with shorter intervals (red line), whereas a constant ratio between inter-trial interval (ITI) and trial duration produces a constant rate of acquisition (green line). (B) Collated data from multiple experiments showing timescale invariance. The median reinforcement to acquisition depends on the ratio between the ITI and the trial duration (from Balsam, Drew, & Gallistel, 2010; permission to reprint pending).

Figure 8 shows simulation results from the replay model and related empirical results from experiments that varied the relative and absolute durations of the ITI. Because of the additional replay iterations during the ITI, in these simulations, we lowered the learning-rate parameter *α* to .01 for original trials and .001 for replayed trials, though the qualitative results are not strongly parameter dependent. Replay speed was tied to trial duration with 10 replays per unit of trial duration during the ITI. As a result, the number of replays per trial depended on both trial duration and the ITI. Acquisition was defined as an associative strength (V) above 0.99. In the replay model, with a constant ITI of 10 units (red line in Fig 8A), the rate of acquisition depends on the trial duration with shorter trial durations producing quicker acquisition because there are more replays of shorter trials during an identical length ITI. With a constant ratio of ITI to trial duration (green line in Fig 8A), there are the same number of replays per ITI and a constant rate of acquisition. These results strongly match the pattern of empirical results (Fig 8B).

A corollary of the principle of timescale invariance is that, with a constant ratio between ITI and trial duration, the absolute number of trials should not influence the rate of conditioning. Indeed, this irrelevance of trial duration has been observed in appetitive conditioning in mice and rats (Gottlieb, 2008, but see Gottlieb & Rescorla, 2010). This observation seems difficult to reconcile with any simple trial-based associative account. Figure 9 (left panel) shows how, in the replay model, the learning from replays during the ITI combined with learning during the trials themselves, can create a balance whereby the number of trials does not influence the speed of learning. With no replays, as in the basic Rescorla-Wagner model (middle panel), learning speed strictly depends on the number of trials per session; whereas with only learning from replays, learning speed depends on the total amount of replay/ITI time, and is therefore inversely related to the number of trials in a session. These two tendencies combine to produce little effect of trial number on acquisition rate. In these simulations, parameters were the same as above, except that session lengths were assumed to last 500 units, each CS lasted 10 units, and the remaining time was devoted to ITI/replays as above. The results in Figure 9 explore the full span from only 1 trial per session and the remaining time serving as one long ITI to 50 trials per session and no ITI/replays.

**Figure 9.**
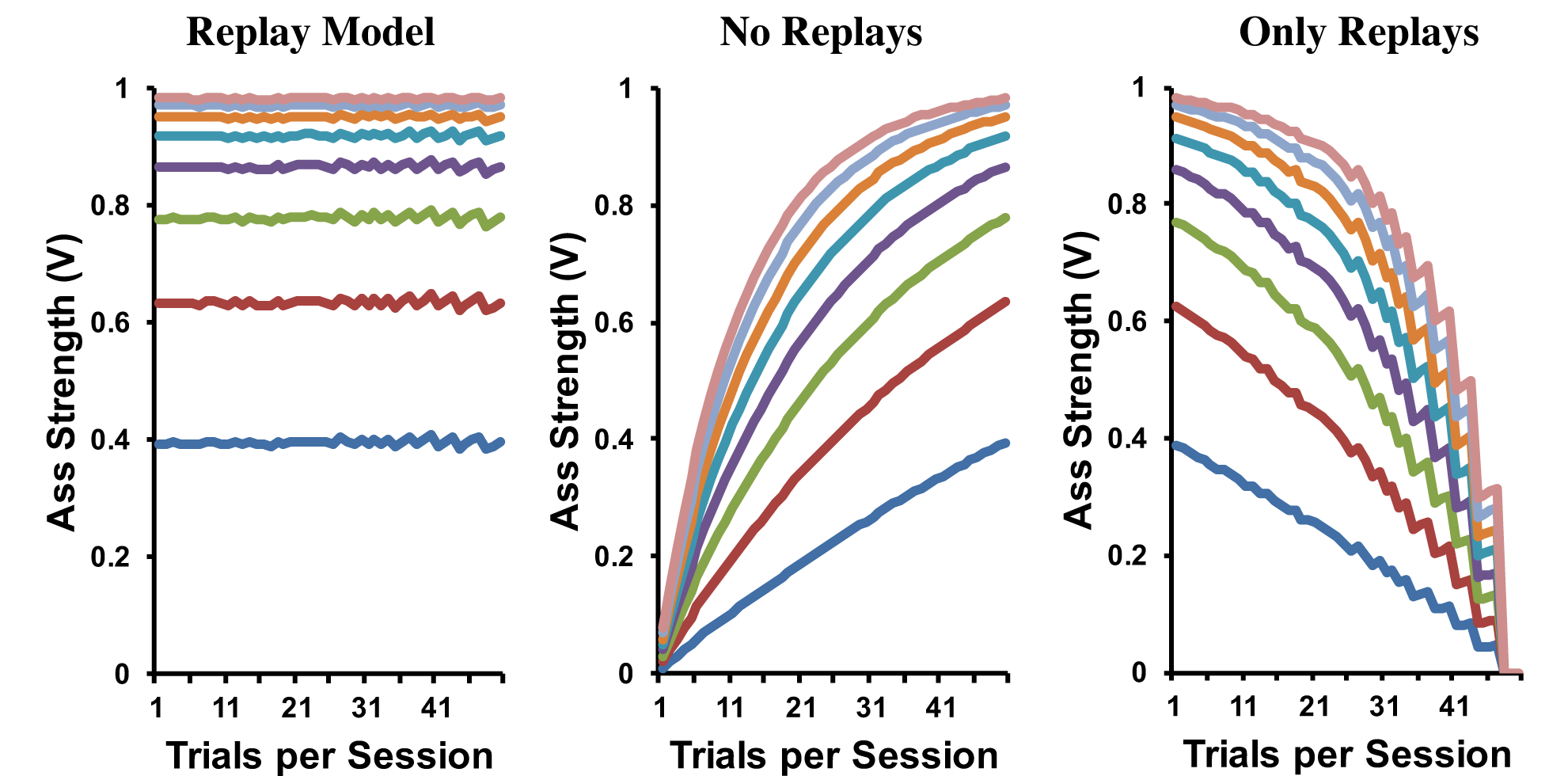
Results from replay model with variable numbers of trials per session. Each successive line is the associative strength at the end of a session (higher lines = later sessions). In the replay model (left panel), the number of trials is irrelevant as the learning from trials balances with learning from replayed experiences during the inter-trial intervals. Without replays (as in the standard Rescorla-Wagner model; middle panel), learning speed increases with the number of trials. With only replays (no learning from experience; right panel), learning speed decreases with the number of trials.

## Selecting Memories to Replay

An important question for the replay model concerns which trials are sampled from the trial memory for subsequent replay. The above simulations all used a simple frequency-based sampling rule, whereby the likelihood of a particular trial type being replayed was proportional to the frequency with which it had previously occurred. Alternatively, one might use a recency-weighted frequency rule, in which the likelihood of selecting a trial for replay depends both on its recency and frequency (e.g., Devenport, 1998). In the experiments discussed above, this altered weighting scheme would have had limited the effects of earlier phases on later conditioning. As long as the recency weighting function is not so steep as to eliminate replay of trials from an earlier phase, the qualitative effects would all be the same, though potentially reduced in magnitude. In spontaneous recovery, for example, so long as some acquisition trials are replayed, then there would be some degree of recovery.

Another possible scheme for selecting which memories to replay involves prioritizing them based on the experienced or expected reward prediction error. From a normative perspective, such a prioritization scheme optimizes the size of the expected update from the replayed trials, focusing on those trials that are least congruent with the current associative strengths. In reinforcement learning, this type of prioritized updating has been shown to enhance the speed with which the *Dyna* algorithm learns (Moore & Atkeson, 1993; Peng & Williams, 1993; Sutton et al., 2008). Indeed, a prioritization scheme suggests why rats in a spatial maze would exhibit experience replay patterns in their hippocampus that tend to return a recently extinguished food location, even after behaviorally the rats have stopped visiting that location (Gupta et al., 2010). For most of the trial-level associative learning experiments we discuss here, replaying based on priority produces very similar results to a frequency-based selection policy. A simple prioritization based on error alone, however, would limit the ability of a replay model to simulate latent inhibition as there were no prediction errors in the first phase and those trials would thereby receive low priority to be replayed.

Tying the likelihood of replay to memory processes also suggests a natural potential extension to this replay model: using contextual similarity to trigger replay. Biasing the replay process towards trials that had occurred in a given context could provide explanations of phenomena such as renewal and the context dependency of latent inhibition and extinction (e.g., Bouton, 1993; Lovibond et al., 1994). As a simple method to implement context dependency, we scaled down the probability of a given trial *T* being selected for replay when the current context was different than the encoding context:

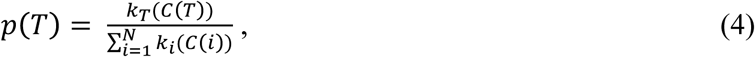

where *C* is the count of the number of trials of a given type in the trial memory (the raw frequency), and *N* is the total number of trial types. For this simulation, *k* was set to 1 for trials that occurred in the same context and 0.1 for trials that occurred in a different context. With only a single context, this formula reduces to simple frequency-based selection, as was used in the earlier simulations. Figure 10 shows how, with this additional assumption, the replay model produces context-depend latent inhibition, whereby the slowing of the learning due to replayed non-rewarded trials occurs more strongly when the context is kept the same. A similar dependence on context would emerge for other phenomena discussed earlier, such as spontaneous recovery, which often exhibits context dependence (e.g., Bouton, 1993).

**Figure 10.**
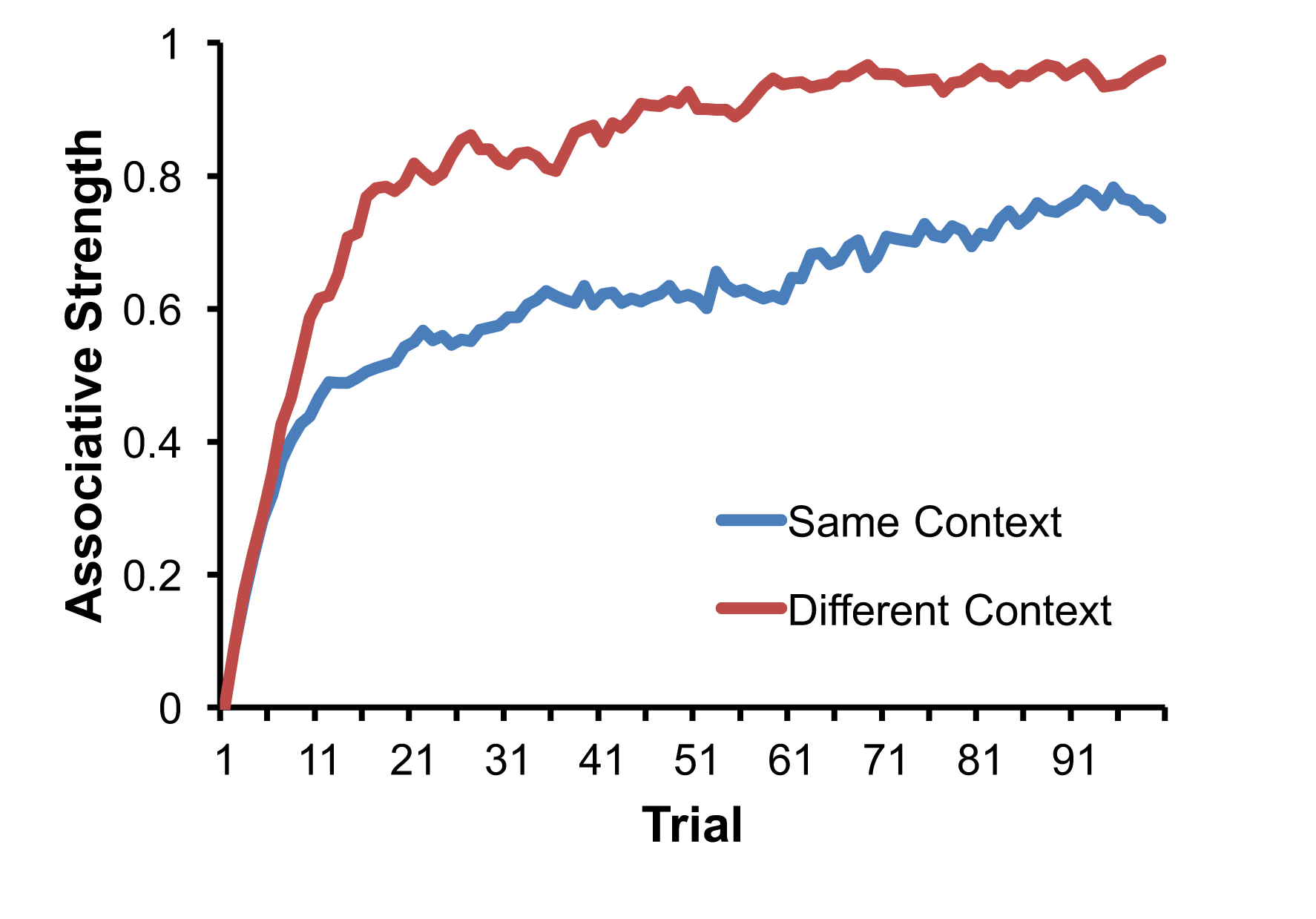
Context-dependent latent inhibition. Associative strength in the replay model as a function of training in the second phase of latent inhibition. One curve (red line) shows when the context has switched between pre-exposure and acquisition, and the second curve (blue line) shows when the same context is used. Acquisition is slower in the same context as non-exposed trials from the first phase are more likely to be retrieved and replayed.

## Further Experiments and Novel Predictions

The replay model also generates many testable predictions about behavior in associative learning experiments. For example, one strong prediction of the model would be that learning can improve in the absence of new experience. The sleep literature contains many examples of such performance enhancements in the absence of further training, including motor skill learning, visual discriminations, and complex cognitive tasks (for a review, see Stickgold, 2005). In the brain, the cortex and hippocampus may interact during sleep leading to the apparent consolidation of learning episodes that occurred while the learner was awake (Stickgold, 2005; Stickgold & Walker, 2007). The replay model suggests a computational mechanism for the positive effects of these interactions, and predicts that these types of improvement should also happen in associative learning tasks, most markedly when the initial training only produces sub-asymptotic behavior. There have indeed also been some associative learning experiments that suggest improved performance in the absence of new training, such as the overnight increases in conditioned responding (e.g., Fig. 1 in Kehoe & White, 2002).

As a further example, consider the experiment of Ohyama and Mauk (2001) that examined the effects of training with different inter-stimulus intervals (ISIs) between the stimulus and reinforcer in classical rabbit eyeblink conditioning. Rabbits were initially given limited training with long ISIs, stopping before the rabbits showed any conditioned responding. The rabbits were then trained with a short ISI until a robust conditioned response was elicited. When rabbits were then presented with a long probe stimulus, they showed two peaks of responding (at the short and long ISIs), even without any additional experience with the longer interval. Our replay model suggests that this increase in responding at the long ISI is due to the replay of trials with the longer interval during the short-interval training rather than any direct effect of the short interval on responding to the longer one. This analysis makes a testable prediction: The mere passage of time (as in spontaneous recovery; see Fig 2) might suffice for an increase in the level of conditioned responding to the longer ISI, even without the short ISI training.

A key assumption of the replay model is that only one trial will be retrieved from the trial memory for replay in any one period. This assumption can be tested experimentally by adding distractor trials. According to the replay model, these irrelevant distractor trials and their replays should reduce the rate of acquisition and the magnitude of other effects that depend on replay. (Figures 2-7; see Salafia & Papsdorf, 1968; Wagner et al., 1973). For example, consider an experiment that has only a few presentations of one reinforced stimulus (X+) versus an experiment with two distinctive reinforced stimuli (X+ and Y+). If the stimuli are sufficiently distinct (no generalization), the replay model predicts that there should be greater improvement in the single-stimulus case because all the replay capacity should be dedicated to that single trial type.

## Limitations and Extensions

As discussed thus far, the replay model only uses close to the simplest memory storage and retrieval mechanisms. The memory starts empty, is entirely infallible, nominally infinite, and non-selective, such that every trial is encoded in memory, and every encoded trial is equally likely to be retrieved and replayed. Of course, any real memory would have a limited capacity, introducing the need for the management of the memory through curation and selection. A curation process would create differences in how trials are retained or entered into memory, perhaps through forgetting or by organizing past instances into types or clusters. A selection process would prioritize retrieval from memory, based on relevant features of the past instances, such as their recency, primacy, saliency, or similarity to the current context (as above).

In its simplest form, the replay model makes an apparently problematic prediction: a single acquisition trial should be sufficient to drive learning to asymptote with sufficient time to replay. In some instances, such as one-trial learning in fear conditioning or the acquisition of phobias (e.g., Bevins & Ayres, 1995; Seligman, 1971), this prediction may be valid, but in most instances of classical conditioning, one trial is insufficient to produce appreciable learning, never mind asymptotic levels of performance. A potential answer to this challenge lies in relaxing in assumptions about the memory contents of this simplest model. For example, relaxing the assumption that the model starts with an “empty” memory which is filled by the single acquisition trial would limit the amount of replay. Alternatively, allowing other events in the life of the animals into the trial memory—i.e., not only those from the experimental context to enter into the memory— would also limit the amount of replay of that single trial. Finally, weighting replay by contextual similarity as above could also limit the amount of replay of that single trial, especially if contextual drift is introduced, whereby as time goes by the context grows more and more different (e.g., Cai et al., 2016; Estes, 1955; Howard & Kahana, 2002),

A further assumption of the replay model is that non-present trials are actively reprocessed. As such, the model supposes a kind of load on the animal’s mental processes that may interfere with the performance of other cognitive tasks or, conversely, may itself be interfered with by the need to perform other tasks (e.g., Pashler, 1994). In particular, replaying past trials would likely place demands on working memory and perhaps on the skilled use of sensorimotor-related mental processes. This connection leads to several natural testable predictions. First, there is the clear prediction that imposing an additional cognitive load, through a secondary or alternative task, would interfere with replay effects. Specifically, the replay model predicts that spontaneous recovery should be diminished through distraction in the interim period, and timescale invariance could be disrupted through the introduction of a secondary task during the inter-trial intervals. Finally, the effectiveness of replay should correlate (on a subject-by-subject or even trial-by-trial basis) with the measured performance on a working memory task, such as delayed matching-to-sample (cf. Gershman et al., 2014 in humans).

The trial memory in the replay model has a similarity to the mental models presumed in model-based reinforcement learning algorithms, which have been used to model various aspects of conditioning and decision-making (Daw, Niv & Dayan, 2005; Balleine & O’Doherty, 2010; Daw et al., 2011). The mental model supposed in those computational approaches typically contains a full abstract model of the world, including the states, actions, and transition probabilities. The trial memory proposed is indeed as a type of mental model, but a particularly simple one, where only instances of past outcomes are stored and recalled. One key advantage of a full mental model, however, is that a single model transition lumps together many real-world transitions, leading to a big savings in the amount that needs to be kept memory. The trial memory proposed here may provide the rudiments for building up such a full mental model. If the memory curation process discussed above could recognize and group trials with similar stimuli and actions, then related transitions could potentially be compactly summarized— perhaps with a transition probability as in the conventional mental models. Moreover, sampling from the trial memory would then provide the same long-run statistics in replayed trials as using the full mental model.

The full replay model can thus be viewed as a light-weight version of model-based planning (Sutton & Barto, 1998). Whereas full planning involves considering every possible action and every possible consequence, the replay model only considers a selected subset of past experiences, achieving many of the same outcomes in a more biologically realistic way with fewer assumptions. In addition, the replay model tightly integrates the learning and planning processes. Instead of dividing value-based processing into separable model-free and model-based computations, the replay model integrates the two processes (see also Keramati et al., 2016; Momennajad et al., submitted; Miller, Shenhav, & Ludvig, submitted). This provides a simple, incremental algorithm that approximates the computation of a full model-based planner.

The replay model, as presented here, is based on the Rescorla-Wagner (RW) learning rule, which is a trial-based, error-correction model of classical conditioning. Each aspect of the RW learning rule can be stepped away from while retaining the replay idea. First, an alternative trial-based learning rule could readily be swapped in instead of the RW learning rule, such as those that compare reward to single associative strengths instead of the sum (e.g., Bush & Mosteller, 1951; Mackintosh, 1975; Pearce & Hall, 1980). Given the many challenges still faced by the RW model (e.g., Miller et al., 1996), this alternative may prove to explain an even wider range of phenomena.

Second, the replay model, like the RW model, is trial level. A potentially interesting extension would be to extend the model to real time. In a real-time version, an experience would consist of multiple steps, and the replay process would traverse these steps, instead of replaying complete trials as a whole. Such an extension would allow the replay model to contend with results from experiments where the relative timing of stimuli or responses are important. Learning rules that extend RW into real-time, such as the temporal-difference (TD) algorithm from reinforcement learning, have already been used to explain many basic classical conditioning phenomena (e.g., Ludvig et al., 2008, 2009, 2011; Schultz et al., 1997; Sutton & Barto, 1990). This combination of the real-time TD learning rule with learning from samples of experience generated from a world model already forms the basis of the *Dyna* approach to model-based reinforcement learning (Sutton, 1990), which has recently been used successfully in the challenging domain of computer Go (Silver et al., 2008, 2016). These computational successes suggest the potential power of a real-time learning rule augmented with a mechanism for learning from replayed or imagined experiences. Adapting this approach as a computational model of classical conditioning, however, raises the challenging question of how stimuli should be represented over time (e.g., Ludvig et al., 2008, 2009, 2012) and the detailed mechanisms of real-time replay (see Johnson & Redish, 2005), but the current trial-level replay model provides the outlines for such an endeavor.

Finally, the replay model could be combined with control methods from reinforcement learning, such as Q-learning or SARSA (Sutton & Barto, 1998; Watkins & Dayan, 1992). These control methods involve actions and are also based on temporal-difference learning. With the inclusion of actions, the replay model could be deployed as a potential model for operant conditioning. With such an action-oriented learning rule, the replay model could potentially be used to provide insight into core instrumental learning phenomena such as latent learning (Tolman, 1929), maze detours (Tolman & Honzik, 1930), and goal devaluation (e.g., Dickinson, 1985). These behaviors are often thought to involve model-based methods, but the replay model could provide a simple memory-based alternative.

## Conclusions and Future Directions

We have presented a model of associative learning that extends the original RW model by incorporating a second process whereby earlier trials are replayed and activate the same associative learning rule. This new model provides an elegant, unifying explanation for a variety of associative learning phenomena that, at first glance, seem completely unconnected, including spontaneous recovery, latent inhibition, retrospective revaluation, and timescale invariance. In addition, we suggested some novel, testable predictions based on the projected effects of the replay process on partially learned information.

The replay model broadens associative theory by supposing that learning can occur from both real and imagined experiences. These imagined experiences may be replayed episodes from memory as above or, more generally, constructed from a model of the world (see Barsalou, 1999; Hassabis & Maguire, 2009; Sutton, 1990). Learning incrementally from this type of simulated experience provides a potential associative mechanism for seemingly complex cognitive phenomena, such as insight learning or incubation (e.g., Sio & Ormerod, 2009; Tolman & Honzik, 1930). The repetition of a range of simulated experience with an incremental learning rule could produce these seeming abrupt changes in overt behavior.

The two pathways (direct and indirect) in the replay model (see Fig. 1) mix learning from recent and remote experiences. In this way, the replay model parallels the complementary learning systems approach to cortico-hippocampal interactions in memory (e.g., Kumaran et al., 2016; McClelland et al., 1995), but the replay model focuses on learning simple associative strengths rather than discovering hidden structure. In the replay model, the direct pathway learns associatively and incrementally from ongoing experience, whereas the indirect pathway immediately stores a relatively veridical representation of the trial or episode. With time-varying experience, the information in these two pathways can diverge, with the direct pathway automatically more sensitive to recent outcomes. The replay mechanism provides a means for making the information in these two pathways more consistent.

More generally, the replay model provides a simple mechanism for incrementally reconciling new information with existing knowledge. The original RW model is path independent. The incremental or decremental effect of each new trial on associative strength completely supersedes the previous associative strength. In the replay model, the past can intrude on the present. New information is immediately stored in the trial memory and only integrated incrementally over time with the associative strengths that drive responding in the model. There is still a bias in the associative strength towards the most recently encountered trials because of the interleaving of learning from real and remembered experiences with their differential learning rates. Nevertheless, the resulting associative strength represents an integration of recent and remote experiences.

In this paper, we introduced a simple model that enhances traditional conceptions of associative learning by allowing for learning from remembered and potentially imagined experiences. Within this broader framework, we have explained several empirical phenomena that pose significant challenges to the traditional associative models. We have also outlined how this modeling approach could open up new avenues for extending associative models to more complex cognitive phenomena.

